# *Toxoplasma* effector *Tg*WIP hijacks dendritic cell actin and motility via Nck1/Grb2 and the WAVE complex

**DOI:** 10.1101/2025.06.22.660983

**Authors:** Pavel Morales, Daniel A. Kramer, Caroline de Moraes de Siqueira, Abbigale J. Brown, Lamba Omar Sangaré, Baoyu Chen, Jeroen P.J. Saeij

## Abstract

The intracellular parasite *Toxoplasma gondii* enhances its dissemination to distant organs by hijacking infected leukocytes via a Trojan Horse mechanism. Upon infecting dendritic cells (DCs), *Toxoplasma* induces a hypermigratory phenotype characterized by podosome dissolution and formation of F-actin stress fibers. We previously showed that these cytoskeletal changes depend on the effector protein *Toxoplasma* WAVE complex-interacting protein (*Tg*WIP) secreted from parasites to infected leukocytes. Here, we identify the host adaptor proteins Non-catalytic region of tyrosine kinase adaptor protein 1 and 2 (Nck1/2) and Growth factor receptor-bound protein 2 (Grb2) as direct *Tg*WIP interactors. *Tg*WIP mainly uses two distinct proline-rich regions (PRRs) to interact with Nck1 and Grb2. Mutating these PRRs abrogates *Tg*WIP binding to Nck1 and Grb2 and diminishes podosome dissolution and DC hypermotility. Furthermore, we show that *Tg*WIP directly interacts with the actin nucleation promoting factor WAVE Regulatory Complex (WRC) via a WRC-interacting receptor sequence (WIRS). Disrupting this interaction also influences actin cytoskeletal remodeling and DC hypermotility. Collectively, our data reveal that *Tg*WIP directly interacts with multiple actin regulators, including Nck1, Grb2, and the WRC, to remodel the actin cytoskeleton of the host cells, elucidating a key mechanism that *Toxoplasma* exploits to enhance host cell migration and dissemination.

## Introduction

The actin cytoskeleton is a highly dynamic network that regulates crucial cellular functions, including cell shape, motility, and internal trafficking (Fletcher and Mullins 2010). Intracellular pathogens have evolved numerous ways to exploit this network to promote their survival, replication, and dissemination. A common strategy is the secretion of virulence factors that modulate host regulators of actin dynamics (Colonne, Winchell, and Voth 2016). Some of these factors use host-mimicry motifs to engage cytoskeletal regulators, inducing actin rearrangements to dampen immune responses or facilitate pathogen spread within and between host cells (Jeng et al. 2004; Chong et al. 2009; Gruenheid et al. 2001). Identifying the molecular mechanisms by which microbial effectors hijack the host cytoskeleton can provide insights into how pathogens manipulate host cells during infection.

*Toxoplasma gondii* is an obligate intracellular pathogen capable of infecting a wide range of warm-blooded vertebrates, including humans and rodents. After oral infection, *Toxoplasma* disseminates broadly in the organism to reach peripheral organs, including the central nervous system (Luft and Remington 1992). While chronic infection is mainly asymptomatic, acute or reactivated infection can cause life-threatening disease in the developing fetus and in immunocompromised individuals (Montoya and Liesenfeld 2004; Schlüter and Barragan 2019).

*Toxoplasma* actively invades peripheral tissues by hijacking leukocytes such as macrophages and dendritic cells (DCs) (Holliman 1989; Channon, Seguin, and Kasper 2000). Infected DCs rapidly undergo a hypermigratory phenotype characterized by cytoskeletal rearrangements and dramatically enhanced motility (hypermotility) (Weidner et al. 2013). *Toxoplasma* induces similar hypermotility features in otherwise immobile macrophages, potentiating parasite dissemination (Ten Hoeve et al. 2022). Infected DCs undergo dissolution of podosomes—actin-rich adhesive structures that degrade the extracellular matrix (ECM) via matrix metalloproteinase secretion—and assemble F-actin stress fibers that promote contractility and motility (Sangaré et al. 2019; Morales et al. 2024; Ólafsson, Varas-Godoy, and Barragan 2018). These cytoskeletal changes have been linked to enhanced systemic dissemination in mice via a Trojan Horse mechanism, whereby infected leukocytes serve as vehicles for parasite transport and immune escape (Lambert et al. 2006; Lambert and Barragan 2010); (Courret et al. 2006).

Although the molecular pathways underlying *Toxoplasma*-induced DC hypermotility remain incompletely defined, they clearly depend on secreted parasite effectors. For example, the *Tg*14-3-3 effector induces hypermotility in both DCs and microglia (Weidner et al. 2016), while the secreted protein kinase ROP17 enhances monocyte locomotion on endothelial surfaces (Drewry et al. 2019). Another key effector is the *Toxoplasma gondii* WAVE-interacting protein (*Tg*WIP), which enhances leukocyte motility independently of toll-like receptor signaling and chemotactic cues (Sangaré et al. 2019; Morales et al. 2024; Weidner et al. 2013). *Tg*WIP is a rhoptry protein secreted into the host cytosol, where it drives podosome dissolution and the hypermotility phenotype of infected DCs (Sangaré et al. 2019; Morales et al. 2024). Mass spectrometry analysis revealed that *Tg*WIP associates with multiple host actin regulators, including Src-homology 2 domain (SH2)-containing phosphatase 1 and 2 (Shp1/2), Non-catalytic region of tyrosine kinase adaptor protein 1 and 2 (Nck1/2), Growth factor receptor-bound protein 2 (Grb2), and components of the WAVE regulatory complex (WRC) (Sangaré et al. 2019). We previously showed that *Tg*WIP is phosphorylated by host Src kinases at Y150 and Y199, which subsequently bind to and activate Shp1/2 (Morales et al. 2024). Whether *Tg*WIP directly interacts with Nck1/2 or WRC remains unknown.

Nck1 and Grb2 are key SH2/SH3 adaptor proteins that couple factors containing phosphorylated tyrosines via their SH2 domains or proline-rich regions (PRRs) via their SH3 domains to actin remodeling pathways (Lettau, Pieper, and Janssen 2009; Lowenstein et al. 1992). Through these interactions, they modulate the activity of Wiskott-Aldrich Syndrome (WAS) family proteins, including WASP and N-WASP, to promote Arp2/3-mediated actin polymerization essential for podosome formation and cell adhesion (W. Li, Fan, and Woodley 2001; Chaki, Barhoumi, and Rivera 2019; Yamaguchi et al. 2005; Aspenström, Lindberg, and Hall 1996). Several pathogens, including enteropathogenic *Escherichia coli* (EPEC) and vaccinia virus, hijack Nck1- and Grb2-mediated actin remodeling to promote infection and spread (Gruenheid et al. 2001; Basant and Way 2022; Ward and Moss 2004). WAVE proteins (WAVE1-3) also drive Arp2/3-dependent actin polymerization essential for cell adhesion and migration (Pollitt and Insall 2009; Takenawa and Suetsugu 2007). WAVE functions within the heteropentameric WAVE regulatory complex (WRC), which contains four other subunits, Sra1, Nap1, Abi2, and HSPC300 (or their corresponding paralog proteins) (Eden et al. 2002; Chen, Chen, et al. 2014a). Various transmembrane receptors recruit the WRC using a conserved WRC-interacting receptor sequence (WIRS) motif that binds to a conserved surface pocket formed by Sra1 and Abi2 (Chen, Chen, et al. 2014a; Rottner, Stradal, and Chen 2021).

We previously showed that *Tg*WIP-Shp1/2 interactions drive F-actin stress fiber-mediated hypermotility in DCs (Morales et al. 2024). However, podosome dissolution occurs independently of this Shp1/2 pathway, suggesting additional mechanisms are involved. Here, we demonstrate that *Tg*WIP’s PRRs directly bind to Nck1 and Grb2, while its WIRS motif binds to the WRC. DCs infected with *Toxoplasma* expressing *Tg*WIP mutants lacking PRR or WIRS sequences fail to exhibit cytoskeletal remodeling or hypermotility. These results reveal a previously unrecognized mechanism through which *Tg*WIP remodels the DC actin cytoskeleton.

## Results

### *Tg*WIP uses distinct PRRs to bind to Nck1 and Grb2

Although *Tg*WIP-Shp1/2 interactions promote the hypermotility phenotype, they do not explain podosome dissolution, suggesting the presence of independent mechanisms. In previous proteomic analysis, Nck1 and Nck2 (Nck1/2) and Grb2 were shown to be potential *Tg*WIP interactors (Sangaré et al. 2019). Given the importance of Nck1/2 and Grb2 in actin regulation and podosome formation, we hypothesized that *Tg*WIP uses its PRRs to interact with the SH3 domains of Nck1/2 and Grb2 (**Fig. 1A**).

**Figure 1.**
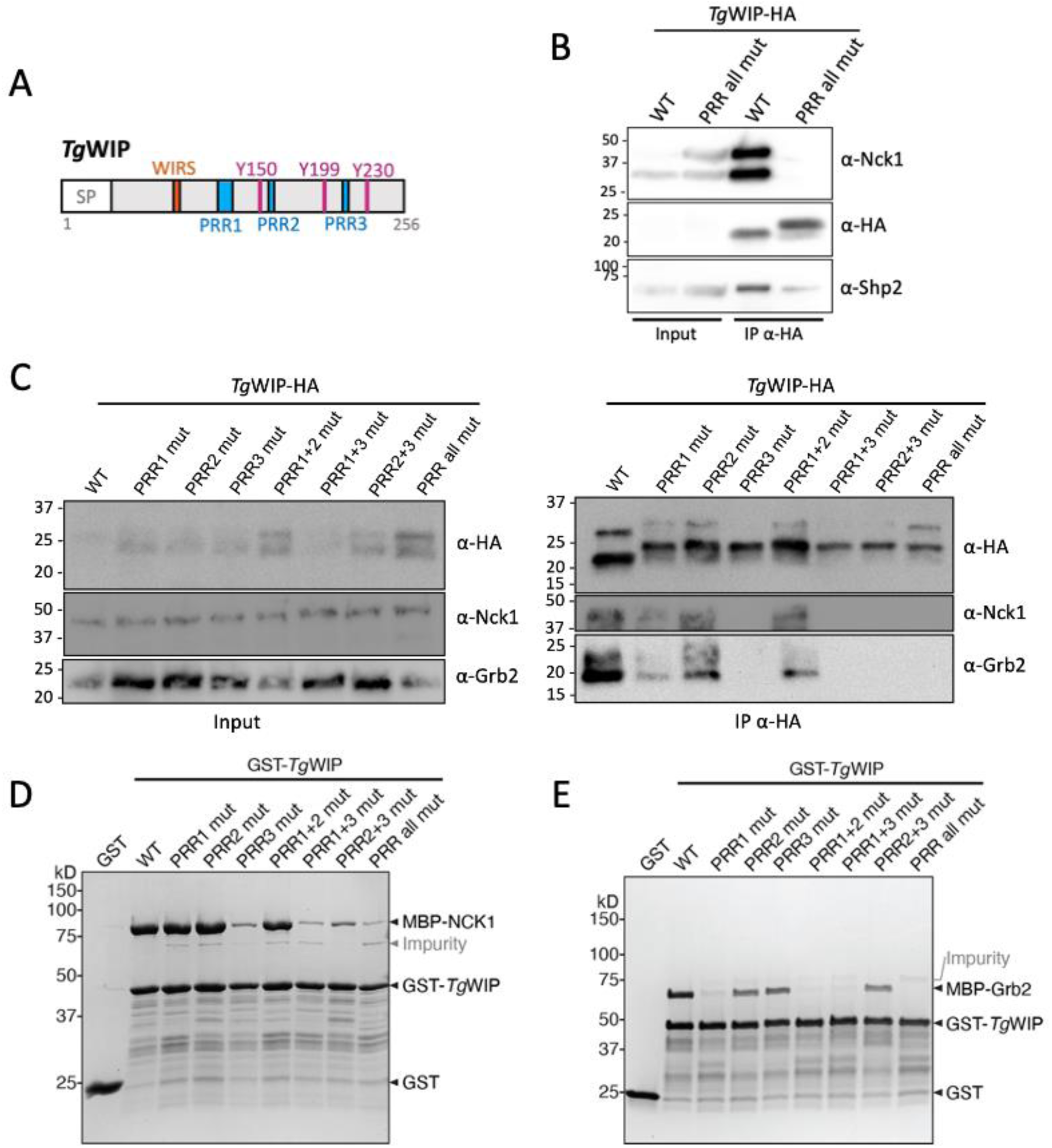
*Tg*WIP uses PRRs to interact with Nck1 and Grb2. (**A)** Schematic showing the amino acid sequence of *Tg*WIP. SP, signal peptide critical for secretion of *Tg*WIP, is excluded from the recombinant protein sequence. (**B)** The murine DC (DC2.4) cell line was infected for 4 h with ME49 Δ*tgwip* parasites complemented with HA-tagged *Tg*WIP^WT^ or *Tg*WIP^PRR^ ^all^ ^mut^. Total cell lysates (Input) and anti-HA immunoprecipitates (IP α-HA) were sequentially immunoblotted with antibodies to Nck1 and HA after stripping the membrane between probes. (**C)** DC2.4 cells were infected as in (B) using ME49 Δ*tgwip* parasites complemented with indicated *Tg*WIP variants. Total cell lysates (Input) and anti-HA immunoprecipitates (IP α-HA) were blotted and reprobed for Nck1, HA, and Grb2, as in (B). (**D-E)** Coomassie blue-stained SDS-PAGE gels showing various GST-tagged *Tg*WIP constructs pulling down MBP-tagged full-length Nck1 in (D) and Grb2 in (E). Each reaction contained 200 pmol of bait and 1000 pmol of prey in a pull-down buffer containing 50 or 100 mM NaCl. Representative gels of two independent experiments are shown.

To test whether *Tg*WIP’s PRRs bind to Nck1, we infected murine DCs (cell line DC2.4) with *Tg*WIP knockout parasites (Δ*tgwip)* complemented with either the wild-type *Tg*WIP with a C-terminal HA tag (*Tg*WIP^WT^) or a mutant in which all three PRRs are substituted with alanine (*Tg*WIP^PRR^ ^all^ ^mut^, **Fig. S1**). Subsequently, we immunoprecipitated HA-tagged *Tg*WIP^WT^ vs. *Tg*WIP^PPR^ ^all^ ^mut^ from infected DCs and used Western blot to detect co-immunoprecipitated Nck1 and Shp2 (**Fig. 1B**). Our results demonstrate that *Tg*WIP^WT^ had a robust binding to Nck1 and Shp2, while *Tg*WIP^PPR^ ^all^ ^mut^ failed to bind Nck1 but retained Shp2 binding. These findings indicate that *Tg*WIP’s PRRs are necessary for binding to Nck1 but dispensable for Shp2 interaction, consistent with the notion that PRRs and phosphotyrosine motifs mediate distinct interactions with host factors.

To identify which PRRs mediate *Tg*WIP binding to Nck1, we generated Δ*tgwip* parasites complemented with *Tg*WIP variants in which different regions of PRRs are substituted with alanine, including PRR1 (*Tg*WIP^PRR1^ ^mut^), PRR2 (*Tg*WIP^PRR2^ ^mut^), PRR3 (*Tg*WIP^PRR3^ ^mut^), or their pairwise combinations (*Tg*WIP^PRR1+2^ ^mut^, *Tg*WIP^PRR1+3^ ^mut^, *Tg*WIP^PRR2+3^ ^mut^). Immunofluorescence confirmed that like *Tg*WIP^WT^, all mutants localized to rhoptries (**Fig. S2**) and were secreted into the host cytosol at levels comparable to *Tg*WIP^WT^. HA-coimmunoprecipitation (co-IP) and immunoblotting from infected DCs showed that *Tg*WIP^WT^ and *Tg*WIP^PRR2^ ^mut^ retained detectable binding to Nck1 and Grb2. In contrast, *Tg*WIP^PRR3^ ^mut^, *Tg*WIP^PRR1+3^ ^mut^, *Tg*WIP^PRR2+3^ ^mut^, and *Tg*WIP^PRR^ ^all^ ^mut^ showed a complete loss of interaction with Nck1 and Grb2, indicating that *Tg*WIP’s PRR3 is crucial for binding to both proteins(**Fig. 1C**). Mutating PRR1 (*Tg*WIP^PRR1^ ^mut^ and *Tg*WIP^PRR1+2^ ^mut^) led to a reduction, but not complete loss of binding to both Nck1 and Grb2, suggesting that PRR1 contributes to, but is not solely responsible for these interactions.

To corroborate the co-IP data and determine whether *Tg*WIP directly binds to Nck1 and Grb2, we performed in vitro GST pull-down assays using purified recombinant proteins. Consistent with the co-IP data, we found that GST-tagged *Tg*WIP^WT^ robustly pulled down MBP-tagged full-length human Nck1 and Grb2, whereas the *Tg*WIP^PRR^ ^all^ ^mut^ had greatly reduced binding to either protein, confirming that the interactions are PRR-dependent (**Fig. 1D,E**). Consistent with the co-IP results, *Tg*WIP^PRR3^ ^mut^ and the PRR3-containing double mutants (*Tg*WIP^PRR1+3^ ^mut^ and *Tg*WIP^PRR2+3^ ^mut^) failed to bind Nck1, identifying PRR3 as essential for this interaction. In contrast, *Tg*WIP^PRR1^ ^mut^ retained Nck1 binding, indicating that PRR1 is dispensable for direct Nck1 interaction in vitro (**Fig. 1D**). For Grb2, *Tg*WIP^PRR1^ ^mut^ had a clear loss of binding, while *Tg*WIP^PRR3^ ^mut^ retained binding, indicating that PRR1 is the primary determinant of the direct interaction with Grb2 (**Fig. 1E**). These results suggest that *Tg*WIP binds Nck1 and Grb2 through distinct PRRs, and that the relative importance of PRR1 and PRR3 differs slightly between cellular and in vitro contexts, possibly due to additional host factors or conformational influences present in cells but absent in the reconstituted system. These interactions, confirmed in vitro, suggest direct binding. However, the cellular context may involve additional host proteins or multiprotein complexes.

### *Tg*WIP uses a WIRS motif to bind the WRC

Previous mass spectrometry data identified all subunits of the WRC as *Tg*WIP binding partners (Sangaré et al. 2019). *Tg*WIP contains a WIRS motif (FGTFVK, corresponding to ɸ-x-T/S-F-x-x, where ɸ represents bulky hydrophobic residues and x for any amino acids) (**Fig. 1A**). Therefore, we hypothesized that this motif mediates the interaction between *Tg*WIP and WRC. Although WRC subunits are difficult to detect by conventional Western blot following co-IP, interaction with the WRC has been consistently observed by mass spectrometry. To determine whether *Tg*WIP binds the WRC directly, we performed pull-down assays using purified, fully assembled recombinant WRC, following protocols used in the original identification of the WIRS motif in transmembrane proteins (Chen, Chen, et al. 2014b). We found that GST-*Tg*WIP, but not GST alone, robustly bound to the WRC in pull-down assays (**Fig. 2B**). Mutating the WIRS motif with alanines substantially reduced the binding (**Fig. 2B**). In parallel, we divided the *Tg*WIP sequence into 12 fragments and purified each fragment as a GST-tagged protein (**Fig. 2C**). Among them, only the F1 fragment, which contains the WIRS motif, was able to pull down the WRC (**Fig. 2C**). Together, these results confirm that *Tg*WIP directly interacts with the WRC via its WIRS motif. This suggests that while the WIRS motif is necessary for direct binding, additional contacts involving *Tg*WIP’s PRRs, such as with SH3 domains in WRC subunits or associated adaptors, may contribute to WRC recruitment in the cellular context.

**Figure 2.**
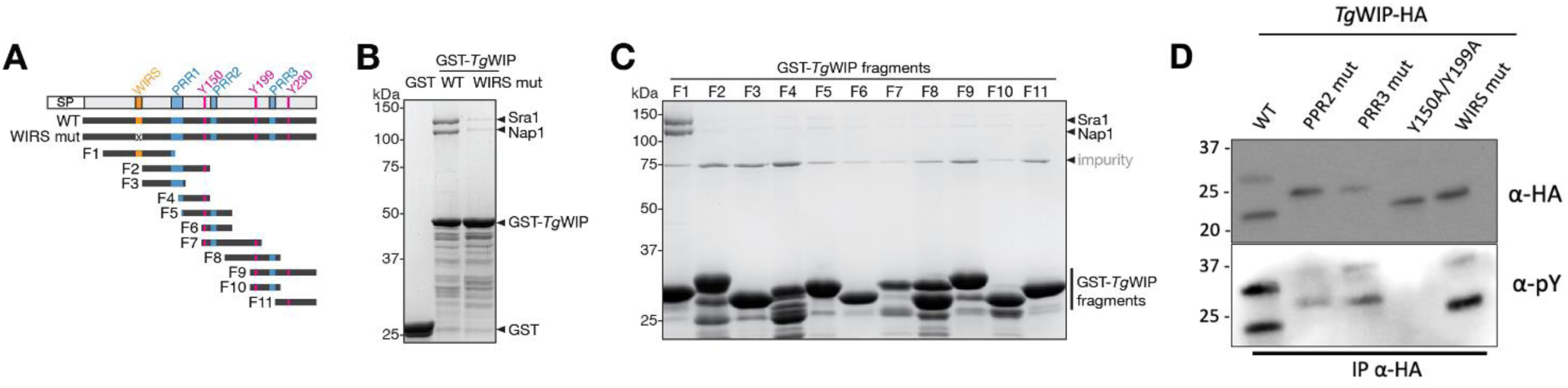
*Tg*WIP engages the WAVE regulatory complex through its WIRS motif while retaining independent Src-mediated tyrosine phosphorylation. (**A**) Schematic showing various purified *Tg*WIP fragments. (**B-C**) Coomassie-blue stained SDS PAGE gels showing GST-*Tg*WIP pulling down recombinantly purified WRC. Each reaction contained 200 pmol of bait and 300 pmol (B) or 60 pmol (C) of prey in a pull-down buffer containing 100 mM (B) or 50 mM (C) NaCl. The bands corresponding to WAVE1, Abi2, and HSPC300 subunits of WRC were out of the gel separation range due to their low molecular weights. Representative gels of two independent experiments are shown. (**D)** DC2.4 cells were infected with *Tg*WIP^WT^, *Tg*WIP^PRR2^ ^mut^, *Tg*WIP^PRR3^ ^mut^, *Tg*WIP^Y150A/Y199A^, or *Tg*WIP^WIRS^ ^mut^ parasites and immunoprecipitated. Shown are the Western blots using HA (first membrane-probing antibody) and phosphotyrosine (pY) (second membrane-probing antibody after stripping membrane) antibodies on total lysate or immunoprecipitated *Tg*WIP.

### *Tg*WIP interacts with Nck1/Grb2 and WRC independently of Shp1/2 interaction

Previous studies established that two tyrosine residues (Y150 and Y199) in *Tg*WIP become phosphorylated by host Src-family kinases (Morales et al. 2024). Once phosphorylated, these sites recruit and activate the host tyrosine phosphatases Shp1 and Shp2, subsequently altering the DC actin cytoskeleton and motility. To test whether mutations in *Tg*WIP’s PRRs or WIRS motif affect its phosphorylation status, we immunoprecipitated *Tg*WIP^WT^, *Tg*WIP^PRR2^ ^mut^, *Tg*WIP^PRR3^ ^mut^, *Tg*WIP^Y150A/Y199A^, and *Tg*WIP^WIRS^ ^mut^ from infected DCs and probed each with anti-phospho tyrosine antibodies. As expected, the Y150A/Y199A double mutant lacked phosphotyrosine signal, confirming that these residues are the primary Src target. All other variants showed detectable, although somewhat less intense, phosphotyrosine signals, compared to *Tg*WIP^WT^ (**Fig. 2D)**. Together, these findings suggest that mutating individual PRRs or the WIRS motif does not abolish Src-dependent *Tg*WIP phosphorylation. This supports the idea that *Tg*WIP engages Shp1/2, Nck1/Grb2, and the WAVE complex, via distinct structural motifs.

### Mutations in *Tg*WIP impair its ability to induce actin cytoskeletal rearrangements

We previously showed that *Tg*WIP’s interactions with the tyrosine phosphatases Shp1 and Shp2 via its SH2-binding motifs Y150 and Y199 is required for F-actin stress-fiber formation in bone-marrow derived dendritic cells (BMDCs) (Morales et al. 2024). However, these interactions did not contribute to *Tg*WIP- mediated dissolution of podosomes, indicating that *Tg*WIP dissolves podosomes in BMDCs via interactions with other host proteins. To test whether the interactions with Nck1/Grb2 or WRC contribute to podosome dissolution, we infected BMDCs with Δ*tgwip* parasites complemented with either *Tg*WIP^WT^ or individual PRR or WIRS mutants (**Fig. 3A**). Consistent with a previous report (Morales et al. 2024), BMDCs infected with *Tg*WIP^WT^ parasites exhibited dramatic cytoskeletal and morphological alterations compared to uninfected or Δ*tgwip-*infected cells, exhibiting increased F-actin stress fibers, membrane protrusions, cell area, and nuclear area, and reduced cell roundness (**Fig. 3F-H**). In contrast, DCs infected with the *Tg*WIP^PRR3^ ^mut^ or *Tg*WIP^PRR1^ ^mut^ strains showed podosome dissolution and F-actin stress-fibers levels similar to Δ*tgwip* parasites. To our surprise, *Tg*WIP^PRR2^ ^mut^ also exhibited similar phenotypes, even though PRR2 was apparently not important for the interaction with Nck1 or Grb2 (**Fig. 1**), suggesting that PRR2 may mediate interactions with additional SH3-containing host proteins involved in actin remodeling. Similarly, BMDCs infected with *Tg*WIP^WIRS^ ^mut^ parasites exhibited increased podosomes expression, reduced F-actin stress fiber levels, fewer membrane protrusions, decreased cellular and nuclear area, and greater cell roundness compared to *Tg*WIP^WT^ infected BMDCs (**Fig. 3A-H**). These results indicate that *Tg*WIP’s PRRs and WIRS motif play a role in modulating the actin cytoskeleton of DCs via interactions with Nck1, Grb2, WRC, and likely other host proteins (**Fig. 3A, B**).

**Figure 3.**
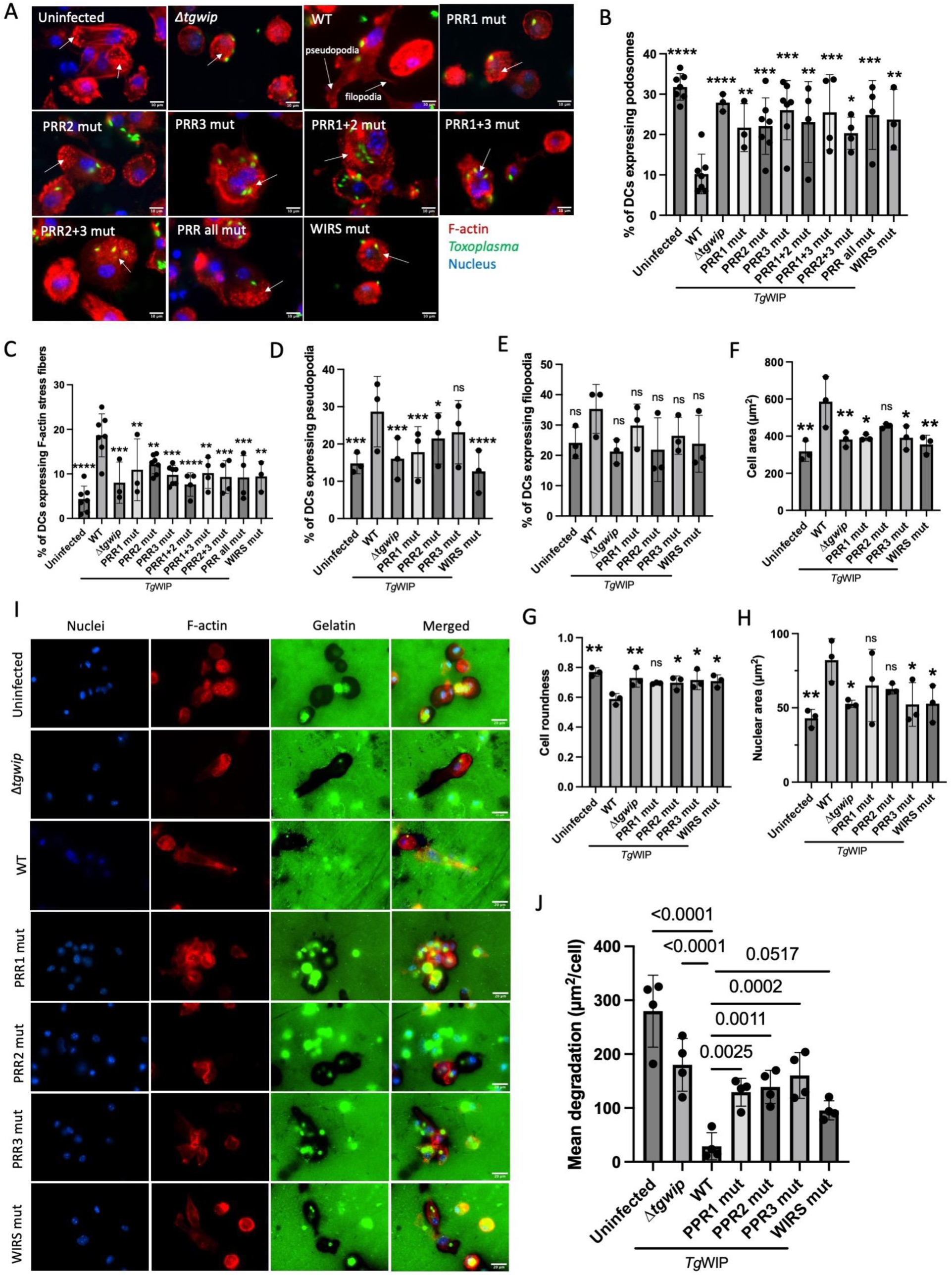
Proline-rich regions and WIRS motif of *Tg*WIP control actin remodeling and matrix degradation in BMDCs. **(A-H)** BMDCs were left uninfected or infected for 4 h with the indicated *Toxoplasma* strains. F-actin and podosomes were visualized with 594 Alexa Fluor Phalloidin, parasites with expressed GFP, and nuclei with DAPI. Unless otherwise indicated, white arrows indicate podosome-expressing, infected DCs. Quantifications are shown at 4 h post-infection of the percentage of BMDCs containing (**B)** podosomes, (**C)** F-actin stress fibers, (**D)** pseudopodia, (**E)** filopodia, (**F)** cell roundness, (**G)** cell area, and (**H)** nuclear area. (**I)** Representative images of pericellular gelatin degradation of BMDCs infected with indicated parasite strains. Loss of fluorescence from FITC-gelatin marks proteolyzed zones. Bright green puncta within the gelatin represent GFP-expressing parasites. F-actin is shown with phalloidin. (**J)** Mean gelatin degradation area per Field of View (FOV) normalized to the cell number (μm^2^/cell) for DCs infected with indicated parasites. Data are from at least three independent experiments. Significance was determined by two-way ANOVA followed by Dunnett’s multiple comparisons. Symbols: (*), (**), (***), (****) indicate statistically significant differences compared to *Tg*WIP^WT^, with p-values of <0.05, <0.005, <0.0002, <0.0001, respectively; ns,not significant.

To further investigate the role of *Tg*WIP’s PRRs and WIRS in dysregulating podosome formation in DCs, we quantified gelatin matrix degradation as a functional readout of podosome activity, a widely used assay that assesses podosome function by quantifying matrix degradation (Ólafsson, Varas-Godoy, and Barragan 2018). DCs infected with Δ*tgwip*, *Tg*WIP^PRR1^ ^mut^, *Tg*WIP^PRR2^ ^mut^, *Tg*WIP^PRR3^ ^mut^ and *Tg*WIP^WIRS^ ^mut^ parasites (**Fig. 3I**) displayed significantly greater gelatin degradation than those infected with *Tg*WIP^WT^ parasites (**Fig. 3J**), indicating impaired podosome dissolution. These results show that both the PRRs and the WIRS motif are individually required for efficient suppression of podosome function and reorganization of the actin cytoskeleton in infected DCs, most likely through interactions with Nck1/2, Grb2, and associated effectors.

### *Tg*WIP mutants could not promote hypermotility

Previous studies have shown that DCs infected with *Toxoplasma* have increased migration across a transwell containing a porous membrane, and this hypermotility phenotype is dependent on *Tg*WIP (Sangaré et al. 2019; Weidner et al. 2013). To test the role of *Tg*WIP PRRs and WIRS motif in modulating the migration of DCs, we performed transwell transmigration assays comparing DCs infected with *Tg*WIP^WT^, *Tg*WIP^WIRS^ ^mut^ and *Tg*WIP^PRR^ ^mut^ strains. Uninfected DCs displayed low levels of transmigration, while priming the cells with lipopolysaccharide (LPS) and adding the DC chemoattractant CCL19 to the bottom chamber of the transwell significantly increased transmigration (**Fig. 4A,B)**. Similar to LPS/CCL19 treatment, infecting DCs with *Tg*WIP^WT^ parasites also significantly increased transmigration, consistent with previous data. In contrast, DCs infected with *Tg*WIP^PRR2^ ^mut^, *Tg*WIP^PRR3^ ^mut^, *Tg*WIP^WIRS^ ^mut^, or Δ*tgwip* parasites displayed significantly lower transmigration compared to *Tg*WIP^WT^ infected DCs. The reduced transmigration was not due to loss of parasites, because the mutations did not affect parasite viability in our plaque assays (**Fig. 4B, C**). Together, our data suggest that *Tg*WIP’s PRRs and WIRS motif are required for elevated transmigration rates in DCs.

**Figure 4.**
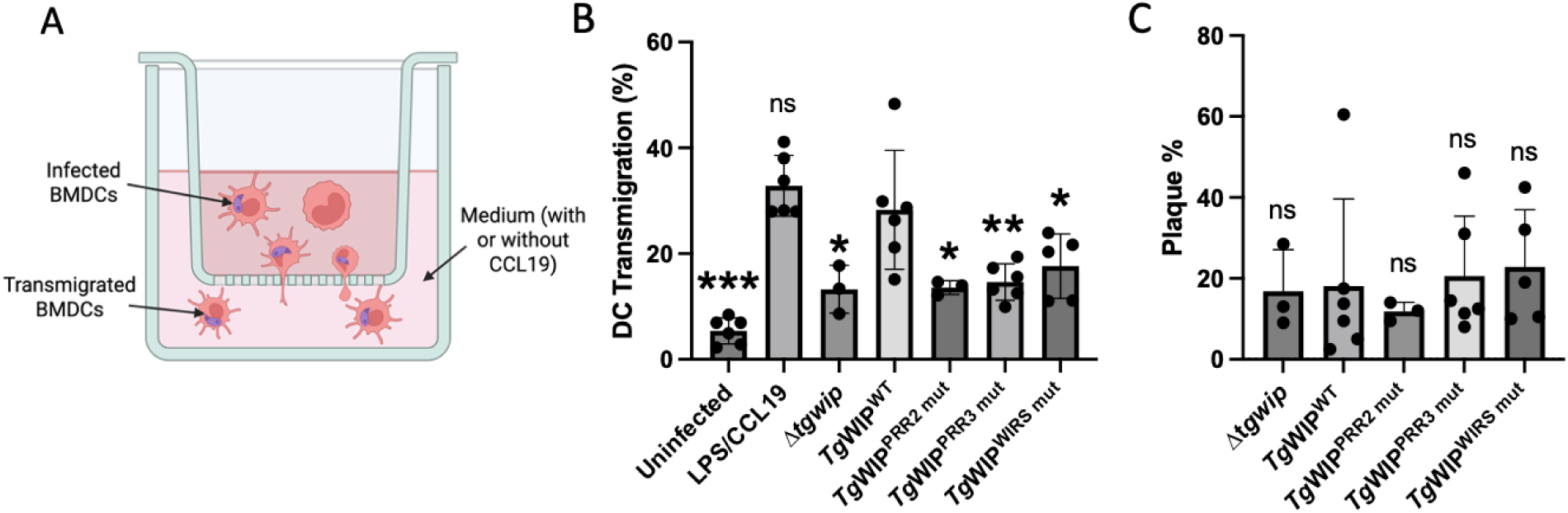
Transwell migration and plaque assays reveal that *Tg*WIP PRRs and WIRS motif promote DC hypermotility. **(A)** Experimental design for the transwell transmigration assay. BMDCs were left uninfected or infected for 4 h with indicated parasites, then loaded in the upper chamber of the transwell. After an 18 h incubation the cells that migrated to the lower chamber were counted. As a positive control, uninfected BMDCs stimulated for 24 h with LPS were allowed to migrate toward the chemokine CCL19. (**B)** Transmigration frequency for each condition expressed as the percentage of total cells added to the insert. Bars show means ±SEM from at least three independent experiments, each performed in duplicate. (**C)** Plaque assay assessing viability of each parasite strain. Bars show mean plaque numbers from the same independent experiments as in (B), each performed in duplicate. Statistics were calculated with two-way ANOVA followed by a Dunnett’s multiple comparisons test. Symbols: (*), (**), (****) indicate statistically significant differences compared to *Tg*WIP^WT^, with p-values of <0.05, <0.005, <0.0001, respectively; ns, indicates not significant.

### Mass spectrometry analysis reveals additional *Tg*WIP ligands involved in actin cytoskeletal regulation

Our Western blot and biochemical analyses show that *Tg*WIP binding to Nck1 and Grb2 is mediated chiefly by *Tg*WIP’s PRR3 and PRR1, but not PRR2 (**Fig.1**). However, DCs infected with *Tg*WIP^PRR2^ ^mut^ also induced actin cytoskeletal rearrangements and low transmigration rates similar to DCs infected with *Tg*WIP^PRR3^ and *Δtgwip* (**Fig. 3, 4)**, indicating the involvement of other host proteins that could modulate actin dynamics in DCs through PRR2. To identify such factors and validate the dependence on PRR1, PRR3, and WIRS motif for Nck1/Grb2 and the WRC, we performed co-IP and mass spectrometry analysis using DC2.4 infected with *Tg*WIP^WT^, *Tg*WIP^PRR2^ ^mut^, *Tg*WIP^PRR3^ ^mut^, or *Tg*WIP^WIRS^ ^mut^ *Toxoplasma.* We included DCs infected with *Tg*WIP^Y150A/Y199A^ parasites as a control. **Table 1** lists the identified host binding partners, along with their relative binding strength to *Tg*WIP^WT^, represented as negative fold changes in peptide counts.

**Table 1.** Identification of host interaction partners of *Tg*WIP. The DC2.4 cell line was infected with indicated parasites expressing HA-tagged *Tg*WIP at MOI 7. After 4 h of infection, DCs were harvested and lysed. *Tg*WIP was then pulled down using the HA tag. The table represents the host interaction partners identified through mass spectrometry. The values represent negative fold changes in peptide counts for each host protein identified in *Tg*WIP mutant protein immunoprecipitations relative to *Tg*WIP^WT^. Components of the WRC (Cyfip1, Cyfip2, WAVE2, Abi1, Nckap1 and Hspc300/Brick1) are highlighted in orange; Shp1 and Shp2 are highlighted in magenta; and the SH3-containing proteins, including adaptors Nck1/2 and Grb2, are highlighted in blue.

| Category | Protein | TgWIP<sup>PRR2</sup> mut | TgWIP<sup>PRR3</sup> mut | TgWIP<sup>Y150A/Y199A</sup> | TgWIP<sup>WIRS</sup> mut |
| --- | --- | --- | --- | --- | --- |
| SH3-containing proteins | Sorbs1 (Cap) | 14 | 30 | 1 | 2 |
|  | Sh3rf1 | 10 | 6 | 2 | 2 |
|  | Grb2 | 6 | 3 | 4 | 2 |
|  | Nck1 | 1 | 60 | 1 | 1 |
|  | Nck2 | 1 | 3 | 1 | 1 |
| Components of the WRC | Abi1 | 4 | 6 | 2 | 4 |
|  | Brk1 (Hspc300) | 4 | 9 | 1 | 3 |
|  | Cyfp2 | 3 | 5 | 2 | 2 |
|  | Wasf2 (Wave2) | 3 | 15 | 2 | 3 |
|  | Nckap1l (Hem1) | 3 | 4 | 1 | 2 |
|  | Abi3 | 3 | 7 | 1 | 1 |
|  | Cyfp1 | 2 | 5 | 1 | 2 |
| SH2-containing proteins | Ptpn6 (Shp1) | 3 | 3 | 32 | 2 |
|  | Ptpn11 (Shp2) | 3 | 2 | 39 | 2 |
| Novel TgWIP interactors | Pstpip1 | 8 | 5 | 1 | 1 |
|  | Nostrin | 6 | 5 | 1 | 3 |
|  | Git1 | 4 | 11 | 2 | 2 |
|  | Arhgef7 | 4 | 332 | 3 | 2 |
|  | Git2 | 2 | 453 | 3 | 2 |
|  | Arhgef7 | 4 | 8 | 2 | 2 |
|  | Arhgef6 | 3 | 7 | 2 | 2 |
|  | Lrrk1 | 3 | 3 | 1 | 1 |

This analysis revealed previously identified *Tg*WIP interactors, including Shp1, Shp2, Nck1, Nck2, Grb2, and components of the WRC (Cyfip1, Cyfip2, WAVE2, Abi1, Nckap1, and Hspc300/Brick1) (Sangaré et al. 2019). Specifically, we observed a stark decrease in Shp1 and Shp2 binding to *Tg*WIP^Y150A/Y199A^, but no change in binding to other *Tg*WIP mutants, consistent with our previous data showing *Tg*WIP’s Y150 and Y199 as primary binding motifs for Shp1 and Shp2 (Morales et al. 2024). This also confirms that mutations in *Tg*WIP’s PRR do not disturb *Tg*WIP’s ability to become phosphorylated and consequently bind to Shp2 (**Fig. 1B,D**). Notably, compared to *Tg*WIP^WT^, binding to Nck1 was drastically reduced (60 fold decrease) in *Tg*WIP^PRR3^ ^mut^ but not in *Tg*WIP^PRR2^ ^mut^. Additionally, in contrast to our biochemical analysis, *Tg*WIP^PRR2^ ^mut^ showed a reduction in binding to Grb2, suggesting that PRR2, while not essential for Nck1 interactions, may contribute to Grb2 binding. The interaction with WRC was also reduced in PRR2 and PRR3 mutants, suggesting the *Tg*WIP-WRC interaction in vivo may involve additional factors, such as potential interaction of the SH3 domain in Abi1/2/3 with *Tg*WIP’s PRRs or the interaction between WRC and Nck1.

Additionally, new *Tg*WIP interactors were also identified. These include important regulators of the actin cytoskeleton, such as adaptor proteins (SH3P12 and SH3RF1), GTPase-activating proteins (GIT1 and GIT2), and guanine nucleotide exchange factors (ARHGEF7/6) (**Table 1**) (Ribon et al. 1998; Manabe et al. 2002; Frank, Adelstein, and Hansen 2006). These proteins either contain SH3 domains, such as ARHGEF6/7, Sorbin, and SH3RF1, or form a complex with SH3-containing proteins (GIT1/2), suggesting that they could interact with the PRRs of *Tg*WIP. Unlike *Tg*WIP^PRR3^ or *Tg*WIP^Y150A/Y199A^, we did not observe the binding of *Tg*WIP’s interactors to be mediated uniquely by *Tg*WIP^PRR2^ ^mut^. Indeed, *Tg*WIP^PRR2^ ^mut^ only displayed shared interactions with *Tg*WIP^PRR3^ ^mut^, highlighting the dominant role of *Tg*WIP’s PRR3 in mediating *Tg*WIP’s binding partners.

## Discussion

Intracellular pathogens have evolved various mechanisms to hijack the host actin cytoskeleton to promote their invasion, replication, and dissemination. We previously showed that the widespread parasite *Toxoplasma gondii* manipulates the actin cytoskeleton and enhances the motility of infected DCs through the secreted effector *Tg*WIP, which interacts with and activates the host phosphatases Shp1/2 (Morales et al. 2024). In this study, we identify an additional mechanism by which *Tg*WIP manipulates the host cell actin cytoskeleton. We show that *Tg*WIP directly interacts with the host adaptor proteins Nck1 and Grb2, as well as the actin nucleation-promoting factor WRC, through distinct motifs. Specifically, *Tg*WIP’s PRR1 and PRR3 sequences are crucial for the interaction with Grb2 and Nck1, respectively, while its WIRS motif is critical for interacting with the WRC. Mutating any of these motifs abolishes *Tg*WIP-mediated podosome dissolution, F-actin stress fiber formation, and hypermobility of infected BMDCs.

Nck1 and Grb2 are critical regulators of the actin cytoskeleton. They contain an SH2 domain and multiple SH3 domains, which function as scaffolds to recruit diverse effectors containing PRRs and phosphotyrosines. Among their major effectors are the nucleation-promoting factors WASP and N-WASP, which activate Arp2/3-mediated actin polymerization, essential for filopodia formation, podosome formation, and cell migration. The multivalent interactions between Nck1, WASP/N-WASP, and additional factors such as Nephrin are known to drive liquid-liquid phase separation (LLPS), which precisely controls the location and timing of actin assembly (P. Li et al. 2012). Our study raises the intriguing hypothesis that *Tg*WIP mimics the PRRs of WASP/N-WASP, thereby interfering with Nck-WASP/N-WASP LLPS and Arp2/3-dependent actin remodeling through direct competition. Similarly, *Tg*WIP could disrupt the interactions of Nck1 and Grb2 with other host effectors containing PRRs and/or phosphotyrosines, such as Cortactin and Gab1 (Grb2-associated binder 1). Given the promiscuity of SH3-PRR interactions, it remains challenging to precisely define the role of *Tg*WIP in modulating related pathways across different cell types and stages of infection.

Our biochemical analysis revealed differences in *Tg*WIP binding to Nck1 and Grb2 depending on the experiment. IPs from murine DCs identified PRR3 as the dominant motif mediating binding to both Nck1 and Grb2, with PRR1 contributing to a lesser extent. In contrast, GST pull-down assays using recombinant human proteins confirmed PRR3 as the primary domain for Nck1 binding, consistent with the IP data. However, in the case of Grb2, the pull-down assays identified PRR1, not PRR3, as the main mediator of the *Tg*WIP-Grb2 interaction. One possible explanation for this discrepancy is the species differences between the experimental systems: IPs were performed in murine DCs, whereas the in vitro assays used human recombinant proteins. Although the orthologs are highly conserved, small differences in SH3 domain sequences could influence PRR binding specificity. Taken together, these results suggest that PRR3 mediates a robust interaction with Nck1, and while it also contributes to Grb2 binding in vivo, PRR1 is the dominant mediator of the *Tg*WIP and Grb2 interaction in vitro.

The WRC is another key actin nucleation-promoting factor, important for lamellipodia formation, phagocytosis, and cell migration. Acting as a signaling hub between the plasma membrane and the Arp2/3 complex/actin, the WRC interacts directly with various transmembrane proteins, particularly cell-adhesion molecules, containing a conserved WIRS motif. The WIRS motif in *Tg*WIP binds to the WRC robustly, which could compete against WIRS-containing membrane proteins, leading to WRC mislocalization and disruption of WRC-mediated actin assembly. In addition, *Tg*WIP could interact with the WRC through alternative routes, including the potential interaction with the C-terminal SH3 domain of the Abi1/2/3 subunit or through indirect interactions by binding to WRC-associated proteins that contain SH3 domains, such as Nck1. This could explain why, in our mass spectrometry analyses, disruption of *Tg*WIP’s PRRs reduced WRC binding more severely than mutation of the WIRS motif, while in our in vitro pull-down assays, the *Tg*WIP-WRC interaction depended exclusively on the WIRS motif. Together, these seemingly conflicting observations suggest *Tg*WIP hijacks WRC-mediated pathways through multiple, potentially redundant mechanisms.

Supporting this, mass-spectrometry also identified a broader network of SH3-containing proteins that associate with *Tg*WIP, including Sorbin (also known as CAP), SH3RF1, and several GAPs and GEFs involved in actin regulation. Interestingly, although PRR2 mutations had little effect on Nck1 or Grb2 binding, they phenocopied PRR3 mutations in cytoskeletal assays. This suggests that PRR2 recruits one or more of these additional SH3-domain partners. Sorbin is a particularly compelling candidate, as it is a scaffolding protein known to regulate the formation of F-actin stress fibers and focal adhesions (Zhang et al. 2006; Ribon et al. 1998). Its interaction with *Tg*WIP via PRR2 may explain how PRR2 mutation results in cytoskeletal phenotypes similar to those regulated by Nck1/Grb2-binding PRR3, reinforcing the idea that multiple SH3-dependent interactions contribute to *Tg*WIP function.

While the simplest model to explain the mechanism of *Tg*WIP is that it uses distinct sequence motifs to bind to different host proteins, with each interaction modulating a specific aspect of host cell behavior, our findings suggest a more integrated mechanism. We have defined the motifs mediating *Tg*WIP’s interactions with Shp1/2, Nck1/Grb2, and the WRC. However, it is striking that disrupting any single interaction results in highly similar, pleiotropic effects on the host actin cytoskeleton and cell migration. This suggests that *Tg*WIP functions as part of a highly cooperative process, depending on the synergistic actions of multiple host protein interactions. Disruption of one interaction likely weakens the overall activity of *Tg*WIP, diminishing its ability to fully remodel the host cytoskeleton. Such functional interdependence could explain the strong evolutionary conservation of *Tg*WIP’s important sequence motifs across different *Toxoplasma* strains. Future studies testing double PRR/WIRS mutants could clarify whether their functions are synergistic or redundant.

*Tg*WIP is one of several *Toxoplasma* effectors that promote leukocyte hypermotility by targeting distinct host actin regulatory pathways. Notably, *Toxoplasma Tg*14-3-3 and ROP17 have also been linked to hypermigration of infected cells. *Tg*14-3-3 is a cytoplasmic and secreted protein with multiple functions in *Apicomplexa*, and its expression in primary DCs and microglia induces a hypermigratory phenotype (Weidner et al. 2016). Although the exact mechanism remains unclear, *Tg*14-3-3 is hypothesized to modulate the Ras-Raf-Mek-Erk MAPK signaling cascade, a pathway implicated in hypermigration of infected DCs (Ólafsson et al. 2020; Ólafsson and Barragan 2020). This is supported by observations that host 14-3-3 proteins accumulate around the parasitophorous vacuole in infected DCs, suggesting sequestration and modulation of MAPK signaling (Weidner et al. 2016). In contrast, ROP17 is a kinase effector that promotes the migration of infected monocytes by targeting GEFs that regulate Rho-ROCK signaling, a pathway that regulates actin nucleation and contractility (Drewry et al. 2019). The distinct host signaling pathways targeted by *Tg*WIP, *Tg*14-3-3-, and ROP17 underscore the possibility for additive or epistatic interactions among these effectors. Together, they highlight the multilayered strategies *Toxoplasma* utilizes to enhance the migratory capacity of distinct leukocytes cell types and promote parasite dissemination.

In summary, this study defines a multifaceted mechanism by which the *Toxoplasma gondii* effector *Tg*WIP hijacks the host actin cytoskeleton to promote DC hypermotility. We demonstrate how *Tg*WIP utilizes distinct sequence motifs to interact with multiple host proteins, Shp1/2, Nck1/Grb2, and the WAVE regulatory complex. This mechanism allows *Tg*WIP to employ pleiotropic effects on podosome and F-actin stress fiber formation, and cell migration. Our findings place *Tg*WIP alongside other *Toxoplasma* effectors such as *Tg*14-3-3 and ROP17 that also target host cytoskeletal regulators to promote immune cell hypermotility. These effectors highlight parasite virulence strategies in which *Toxoplasma* hijacks host cytoskeleton networks to disseminate efficiently within the host. This work not only builds on the molecular mechanisms used by intracellular pathogens but also reinforces the known host molecular pathways that govern the actin cytoskeleton and cell motility.

## Acknowledgements

We thank Michael Rosen lab at UTSW for sharing NCK1 and Grb2 constructs.

## Author Contributions

J.P.J.S. and B.C. conceived the project. J.P.J.S. oversaw cell biological and proteomic experiments performed by P.M. and D.M.S. B.C. oversaw protein purification and biochemical experiments performed by D.A.K. and A.J.B. P.M drafted the manuscript and prepared the figures with assistance from all other authors.

## Materials and Methods

### Parasite Culture

All the *Toxoplasma* parasite strains were routinely maintained *in vitro* in monolayers of HFF at 37 °C in 5% CO_2_ as previously described (Rosowski et al. 2011).

### Primary Host Cell Culture

Bone marrow-derived Dendritic Cells (BMDCs) were isolated from female CD1 mice (Charles River Laboratories) of 5-to-8 weeks old as previously described (Fuks et al. 2012). BMDCs were obtained by culturing murine bone marrow cells in Roswell Park Memorial Institute (RPMI) 1640 with 5% FBS, 10 mM HEPES (pH 7.9), 50 µM 2-mercaptoethanol, 10 µg/mL gentamicin, referred to as complete medium (CM), and supplemented with recombinant mouse GM-CSF (40 ng/mL, Peprotech) and mouse IL-4 (40 ng/mL, Peprotech). Loosely adherent cells were harvested after 8 days of maturation. The medium was changed every 2 days in culture with fresh GM-CSF and IL-4 (modified from Inaba et al. (Chen, Padrick, et al. 2014)).

### Co-immunoprecipitation

DC2.4 cells were grown in a 150-mm culture dish until 100% confluency and infected (MOI: 5 to 7) for 4 h with ME49 *Tg*WIP^WT^ parasite expressing *Tg*WIP mutant strains or mock infected, using multiplicity of infection (MOI) of 5 to 7. Cells were then scraped in PBS, centrifuged, and resuspended in 1 or 3 mL of lysis buffer (HEPES 10 mM pH 7.9, MgCl_2_ 1.5mM, KCl 10 mM, EDTA 0.1 mM, dithiothereitol (DTT) 0.5 mM, NP40 0.65%, cocktail of protease inhibitor (Roche), phenylmethylsulphonyl fluoride (PMSF) 0.5 mM) for 30 min at 4 °C. The lysate was centrifuged for 30 min at 18,000 x g, 4 °C. Each sample was incubated with 35 μl of magnetic beads coupled with HA antibodies (Thermo scientific) and placed on a rotator overnight at 4 °C. The beads were washed three times with Tris-HCl 10 mM pH7.5, NaCl 150 mM, Triton X-100 0.2%, PMSF 0.5 mM, a cocktail of protease inhibitors (Roche), followed by one-time wash with Tris-HCl 62.5 mM pH6.8, and finally resuspended in 100 µl of this buffer for analyses. Bound proteins were eluted by boiling the samples for 5 min.

### Immunoblotting

For each sample, 30 μl of the HA magnetic beads of each sample was used to run on a 12% SDS-PAGE gel. The proteins were transferred to a PVDF membrane, blocked 30 minutes with Tris-buffered saline with Tween 20 (TBST) containing 5% nonfat dry milk. The membrane was blotted overnight at 4°C with a rat antibody against the HA tag (Sigma Aldrich, Cat#11867431001, add the exact number for all antibodies 1:500 dilution), phosphorylated Tyrosine (Cell Signaling, Cat#8954S, 1:500 dilution), Shp1 (Invitrogen, Cat#3759, 1:1000 dilution), or Shp2 (Cell Signaling, Cat#3752S, 1:1000 dilution) antibodies, followed by their respective secondary antibodies (ThermoFisher Scientific, 1:5000 dilution).

### Proline-Rich Region Mutants Generation

The prolines in the PRR regions were replaced by alanines, either individually or in combination, to create single, double, and triple PRR mutants. The sequences were synthesized by Twist Biosciences and incorporated into the complementation plasmid pUPRT::TgWIP-HA (Sangaré et al, 2019). *Toxoplasma* ME49 Δ*tgwip* were transfected with the complementation plasmid, using Gene Pulser Xcell™ (BioRad), as previously described by Soldati and Boothroyd (1993). The transfected parasites were selected with 10 μM of FUDR (5-fluorodeoxyuracil) and cloned by limiting dilution.

### Podosome Assay

To test the podosome dissolution after *Toxoplasma* challenge, BMDCs were seeded on collagen-coated glass coverslips overnight, after which freshly egressed parasites were added at MOI 3 for 4 h. The coverslips were then washed with PBS, fixed with 4% paraformaldehyde (PFA) for 20 min, permeabilized for 5 min with 0.1% Triton X-100, and blocked for 1 h with PBS with 3% (w/v) BSA (blocking buffer). The coverslips were incubated with DAPI to stain the host nucleus, and with Alexa Fluor 488 Phalloidin in blocking buffer for 30 min to visualize F-actin. GFP-expressing *Toxoplasma* were detected through their intrinsic GFP signal. The coverslips were mounted with Vecta-Shield mounting oil and the microscopy was performed with NIS-Elements software (Nikon) and a digital camera (CoolSNAP EZ; Roper Scientific) connected to an inverted fluorescence microscope (eclipse Ti-S; Nikon) and either phase contrast or DIC imaging. Podosomes were identified and quantified as described in (Weidner et al. 2013).

### DC Gelatin Degradation Assay and Image Analysis

The *in vitro* gelatinolytic activity of DCs was analyzed by gelatinolysis of Oregon green 488 (OG 488)-conjugated porcine gelatin (Molecular probes). DCs (2.5 × 10^4^/well) infected with freshly egressed *Toxoplasma* tachyzoites (ME49-GFP, MOI 3) were deposited on OG-488 gelatin-coated glass coverslips and incubated for 24 h in CM. Cells were subsequently fixed (4% paraformaldehyde, Sigma), stained with DAPI (Invitrogen) or Alexa Fluor 594 Phalloidin (Thermo Fisher). Imaging and analysis were performed as indicated below. The *in vitro* gelatinolytic activity of DCs was analyzed by gelatinolysis of Cy-3-conjugated gelatin (molecular probes). After fixation (4% PFA, Sigma), cells were stained with DAPI (1:1000) and Alexa Fluor 594 Phalloidin (1:800). Using the software ImageJ version 1.54, the threshold of the green channel (Alexa Fluor 488 Phalloidin) was adjusted to distinguish single cells. The tool ‘‘Analyze particles’’ was used to count cells and measure the area of degradation. Gelatin degradation was defined as loss of signal (gelatin, Oregon green-488). The degradation of 100 cells was manually quantified for each condition. The “Analyze particles” tool in ImageJ was used to quantify DC cell area and roundness based on F-actin staining with phalloidin, which outlines the full cell body. The same tool was also used to measure nuclear area using DAPI to visualize nuclei.

### Transmigration Assay

The transmigration assays were performed by culturing BMDCs in CM with the addition of freshly egressed *Toxoplasma* strains (MOI 3) for 4 h. DCs were then transferred to transwell filters consisting of a porous membrane (8 μm; Corning) incubated at 37 °C. After 18 h, DCs from the bottom chamber were collected and quantified by hemocytometer.

### Protein Purification

See Supplementary Table 1-2 for the amino acid sequence of all *Tg*WIP constructs used in the pulldown experiments. All GST-3C-*Tg*WIP PRR mutant constructs were synthesized by ThermoFisher GeneArt and sequence verified independently. All *Tg*WIP constructs, as well as Nck1 and Grb2 constructs, were expressed in Arctic Express (DE3) RIL (Agilent) cells, using induction with 1 mM IPTG at 10°C for 16 hr. Cells were disrupted by sonication, followed by centrifugation at 19,000 rpm at 4 °C for 30 minutes. Clarified supernatant was mixed with GSH-Sepharose beads (Cytiva) for 30 minutes at 4 °C, followed by three washes and elution using a buffer containing 30 mM reduced GSH and 100 mM Tris pH 8.5. Eluted proteins were further purified using Source 15S cation chromatography (Cytiva) at pH 8, followed by size exclusion chromatography using a Superdex 200 Increase gel filtration column (Cytiva). The WRC was purified and assembled as previously described (Chen, Padrick, et al. 2014).

### Pull-down Assay

Pull-down assays were performed as previously described (Kramer et al. 2023). Nck1 and Grb2 experiments were performed with 200 pmol of bait (GST-*Tg*WIP), 1,000 pmol of prey (MBP-Nck1 FL or MBP-Grb2), and in 100PDB buffer (100 mM NaCl, 10 mM HEPES pH 7.0, 5% (wt/vol) glycerol, 5 mM 2-mercaptoethanol, and 0.05% Triton X-100). WRC experiments were performed with 200 pmol of bait (GST-*Tg*WIP) and 60-300 pmol of prey (WRC), and in 100PDB or 50PDB buffer (50 mM NaCl, 10 mM HEPES pH 7.0, 5% (wt/vol) glycerol, 5 mM 2-mercaptoethanol, and 0.05% Triton X-100). In each reaction, 20 µL of GSH-Sepharose beads (Cytiva) were mixed with bait and prey proteins in 1 mL of pull-down buffer. The samples were mixed at 4 °C for 30 min and then washed three times with 1 mL of pull-down buffer. Protein was eluted with 40 µL of elution buffer (containing 30 mM reduced glutathione and 100 mM Tris pH 8.5), and the eluate was examined by SDS-PAGE and Coomassie blue staining. Pull-down experiments were repeated at least twice to ensure reproducibility of the results.

## Supplemental Figures

**Fig S1.**
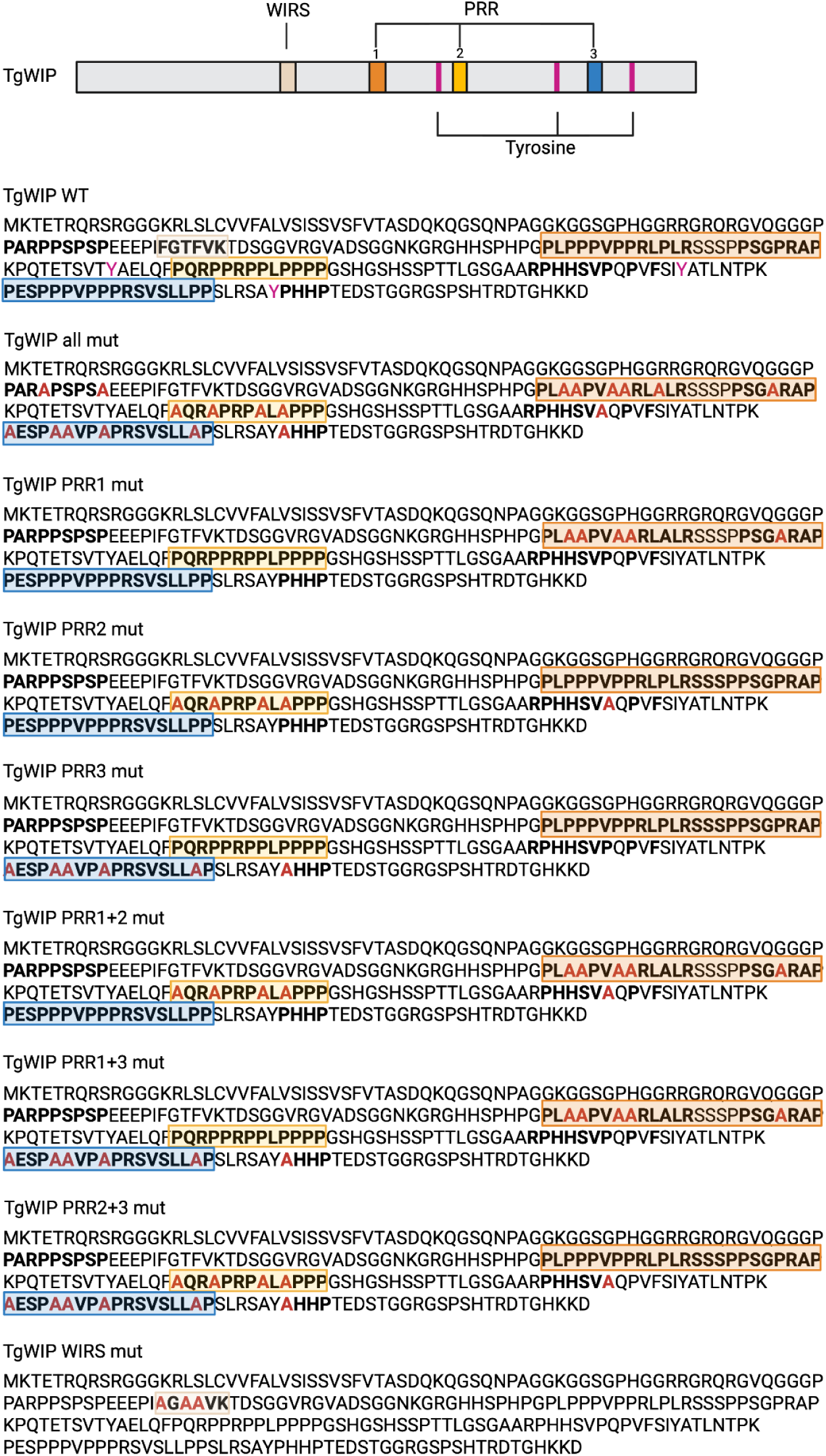
Alignment of *Tg*WIP amino acid sequences and mutant strains. ME49 *Tg*WIP amino acid sequence of strains *Tg*WIP^WT^, *Tg*WIP^PRR^ ^all^ ^mut^, *Tg*WIP^PRR1^ ^mut^, *Tg*WIP^PRR2^ ^mut^, *Tg*WIP^PRR3^ ^mut^, *Tg*WIP^PRR1+2 mut^, *Tg*WIP^PRR1+3^ ^mut^, *Tg*WIP^PRR2+3^ ^mut^, and *Tg*WIP^WIRS^ ^mut^. Mutated residues in each strain are indicated. Motifs are color-coded: PRR1 in orange, PRR2 in yellow, PRR3 in blue, tyrosine residues in magenta, WIRS motif in tan.

**Fig S2.**
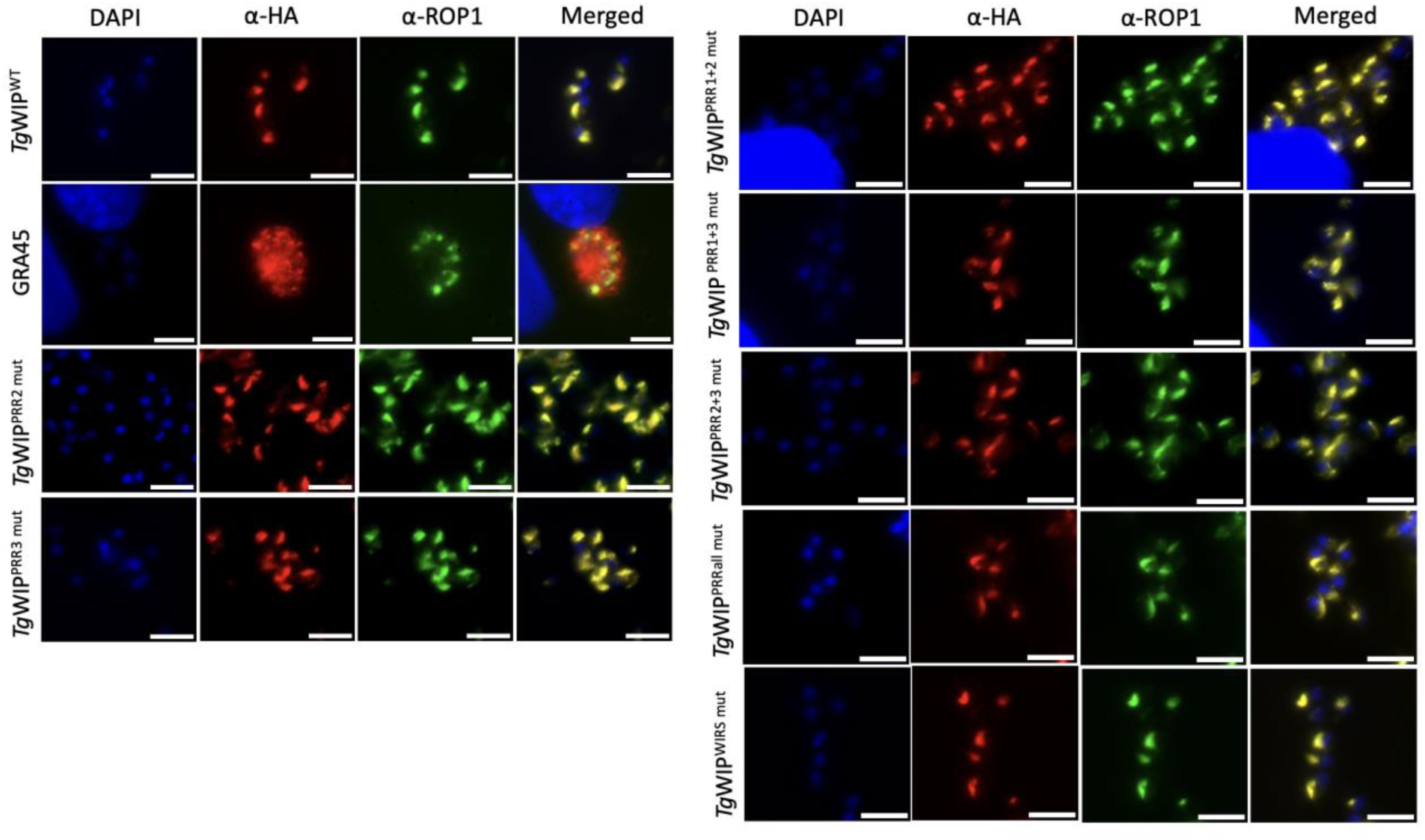
Rhoptry organelle localization of *Tg*WIP mutants. *Tg*WIP and rhoptry organelle colocalization of *Tg*WIP^WT^, *Tg*WIP^WIRS^, or *Tg*WIP^PRR^ mutant strains. Immunofluorescence assays on Human Foreskin Fibroblasts (HFFs) infected with *Toxoplasma* containing either endogenously HA-tagged wildtype *Tg*WIP (*Tg*WIP^WT^) or *Tg*WIP expressing PRR mutations. ROP1 was used as a marker for rhoptry organelles. GRA45 is a *Toxoplasma* secreted protein that localizes in the PV lumen, the GRA45 *Toxoplasma* strain was used as a negative control.

**Fig S3.**
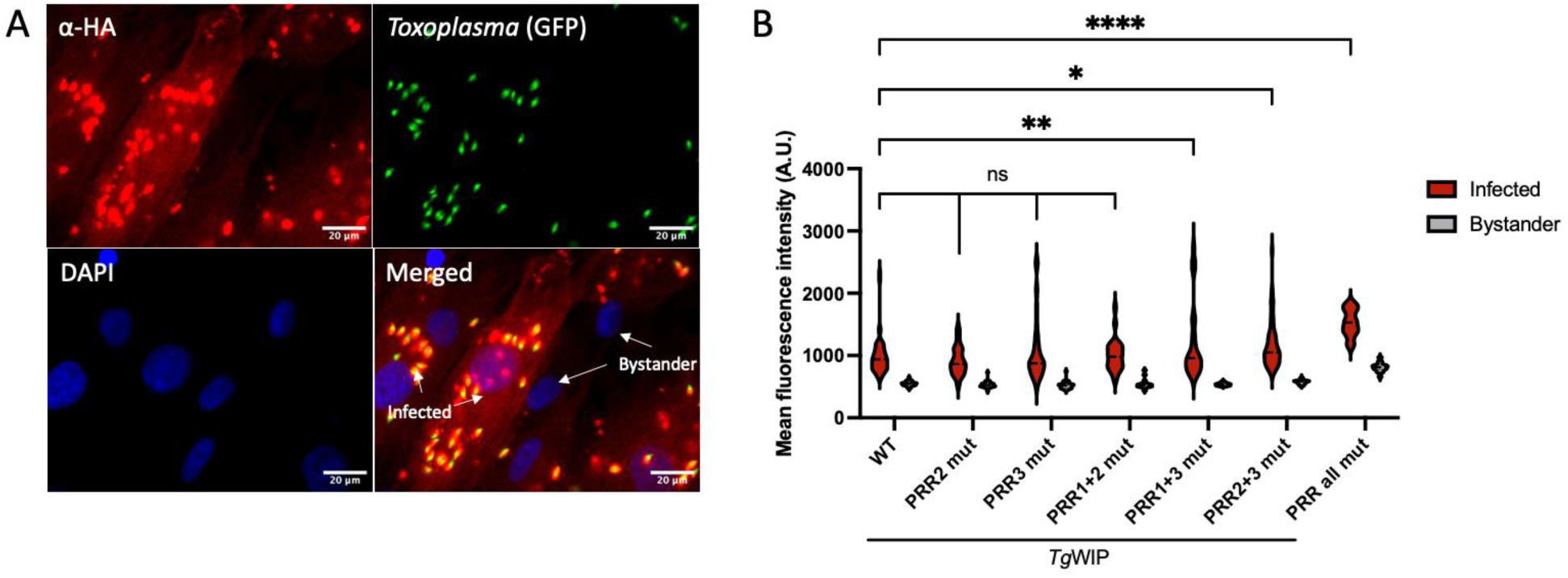
Quantification of *Tg*WIP-HA signal in HFFs infected with *Toxoplasma Tg*WIP^WT^ or *Tg*WIP^PRR^ mutants. **A)** Representative immunofluorescence images of *Tg*WIP^WT^ *Toxoplasma* infected HFFs stained for HA. White arrows indicate uninfected bystander cells. **B)** The mean HA fluorescence intensity of uninfected (bystander) HFFs or HFFs infected with *Toxoplasma Tg*WIP^WT^ or *Tg*WIP expressing PRR mutations was determined after 3 h infection.

**Fig S4.**
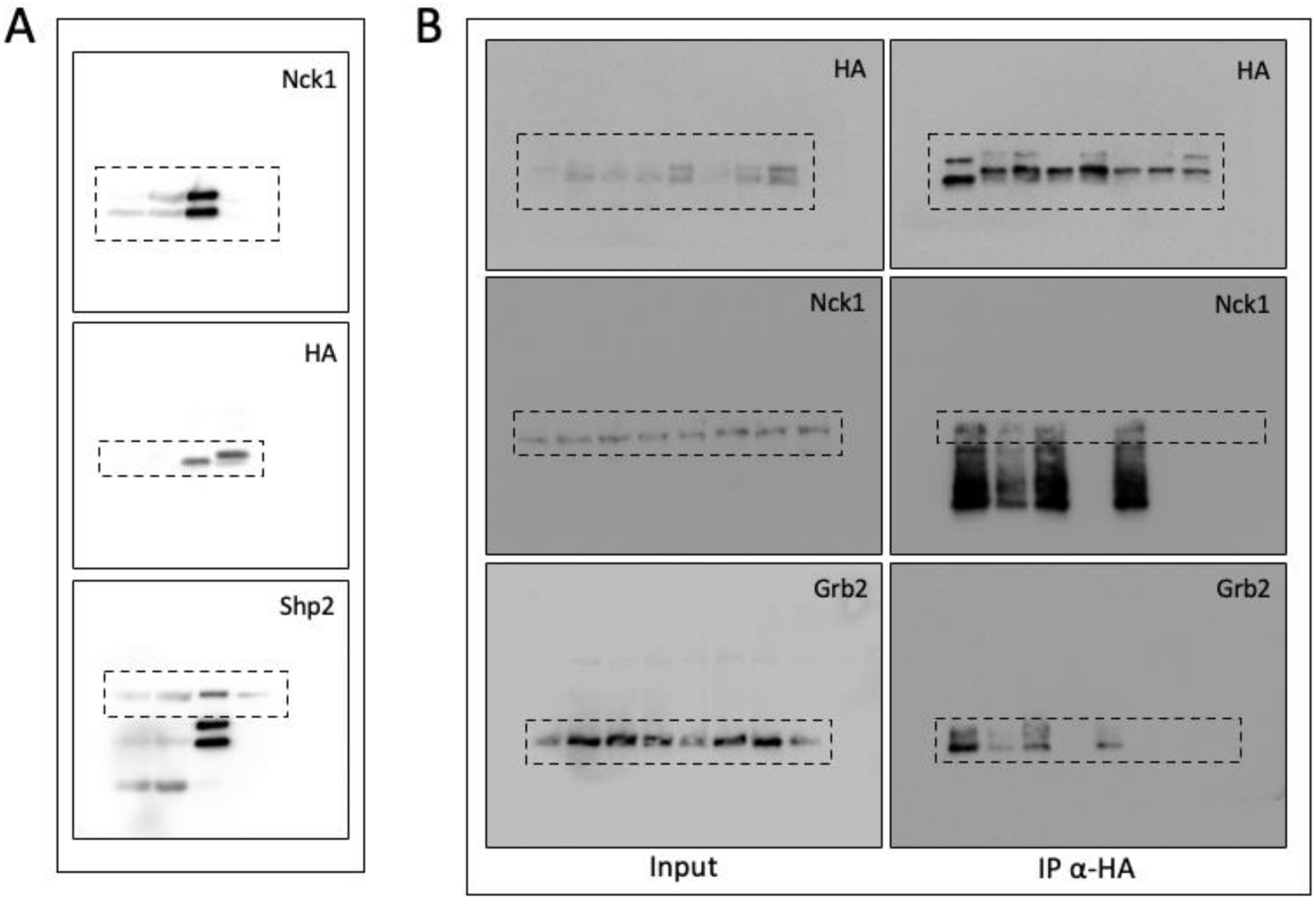
Uncropped Western blot images for main figures. Uncropped Western blot images in this study. **A)** The membrane was sequentially probed with antibodies against Nck1 (first), HA (second, after one stripping step), and Shp2 (third, after a second stripping). **B)** The membrane was first probed with HA antibody, followed by Nck1 (after one stripping step), and Grb2 (after a second stripping).

**Supplementary Table 1.**
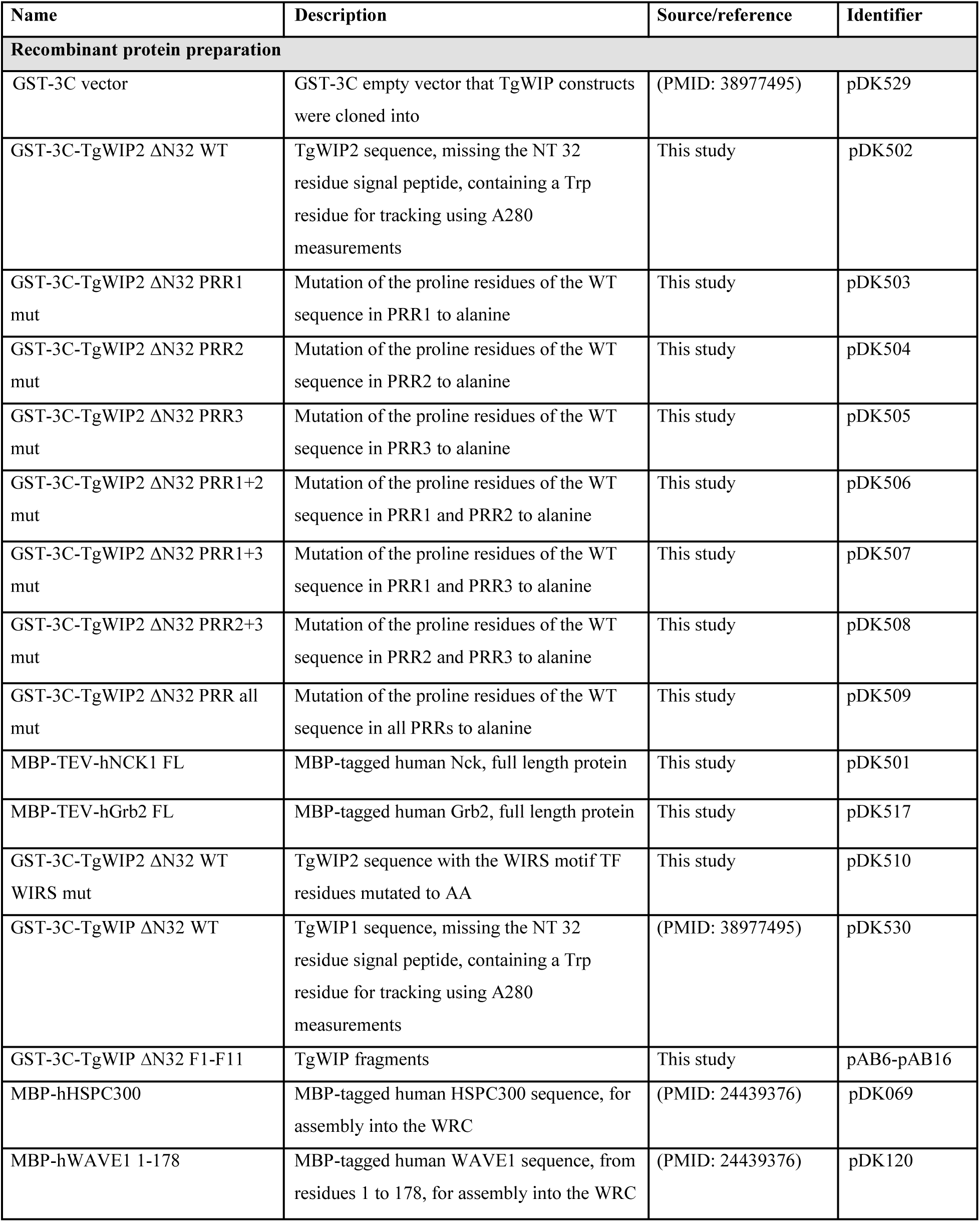
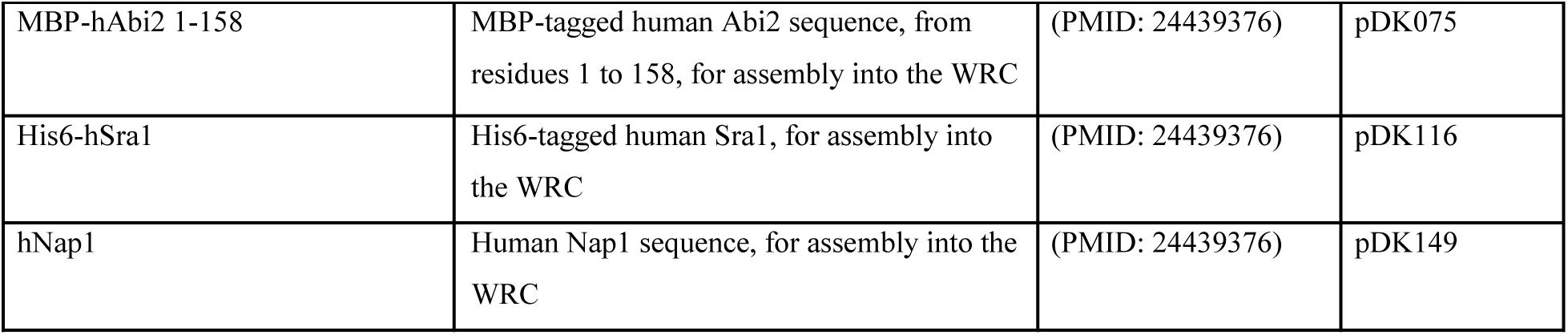
DNA constructs used in this study.

**Supplementary Table 2.**
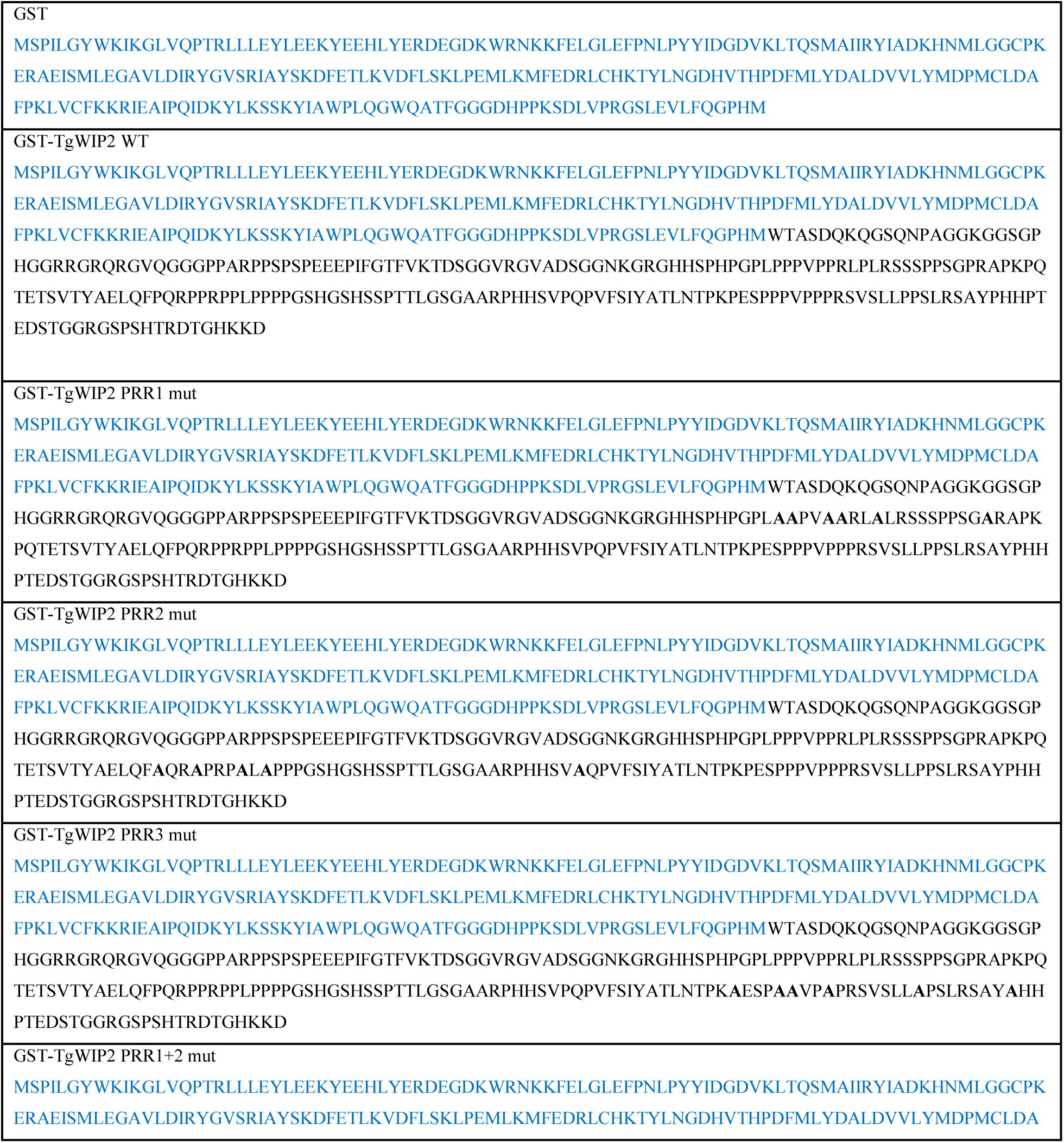
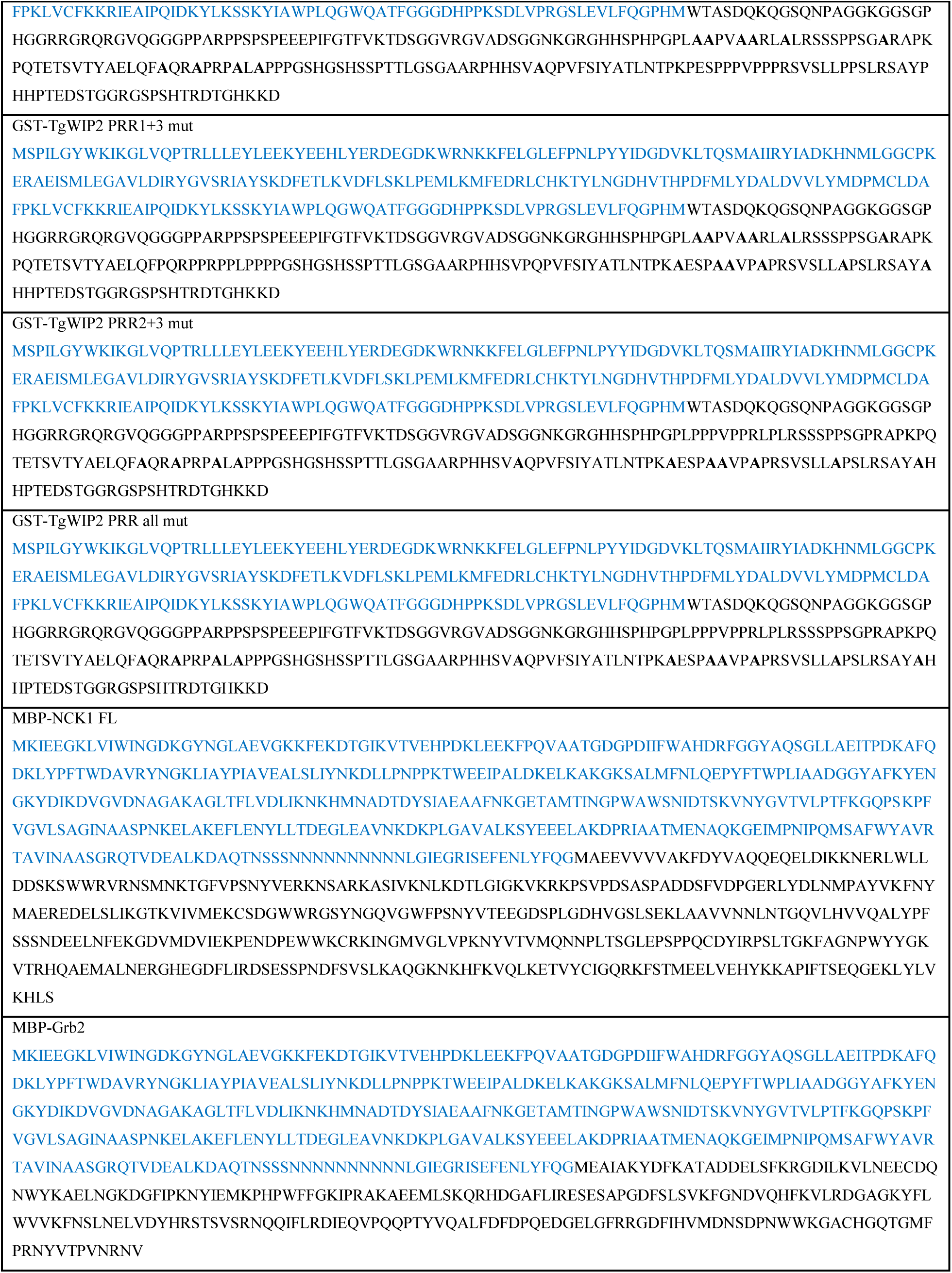
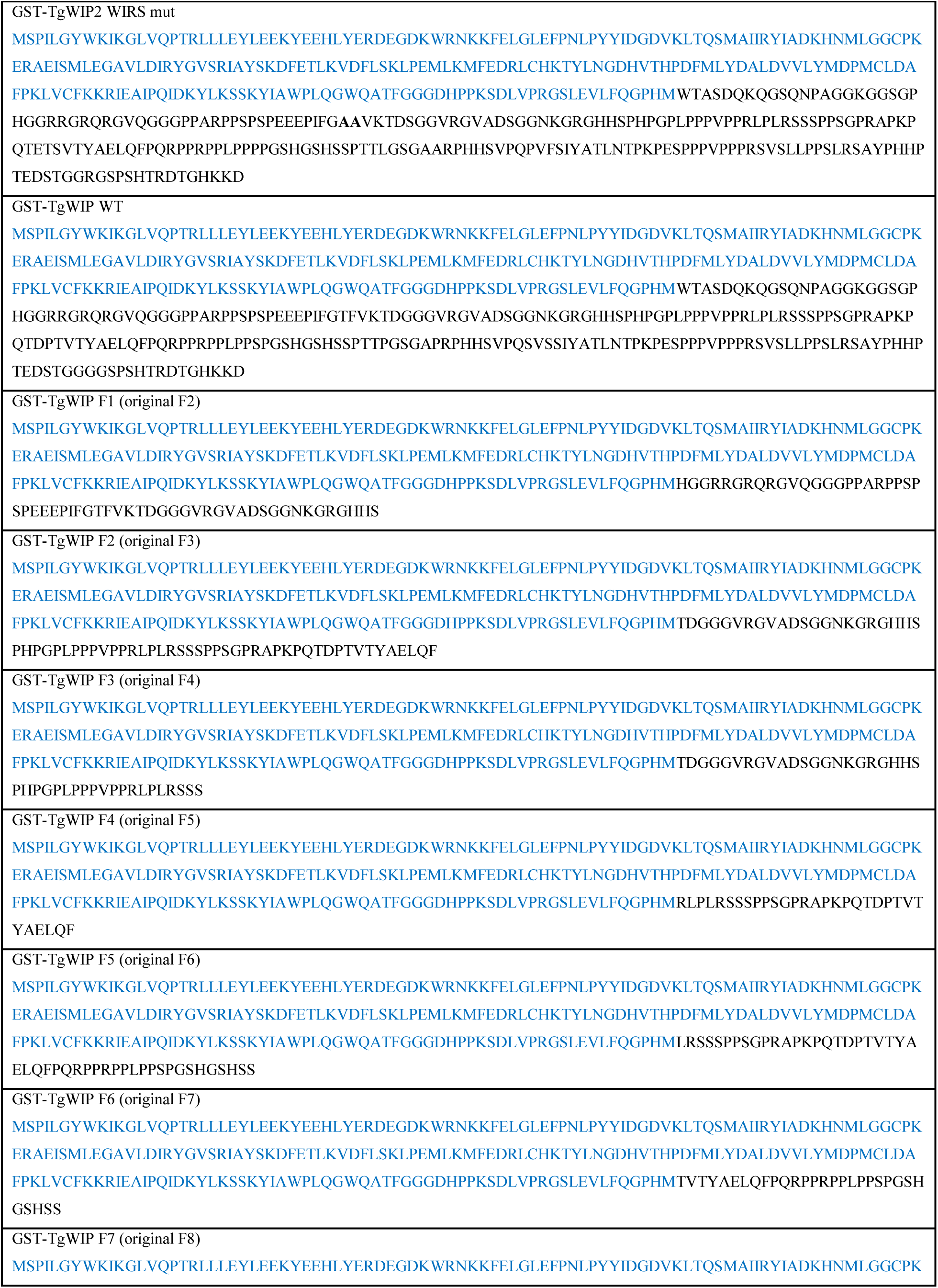
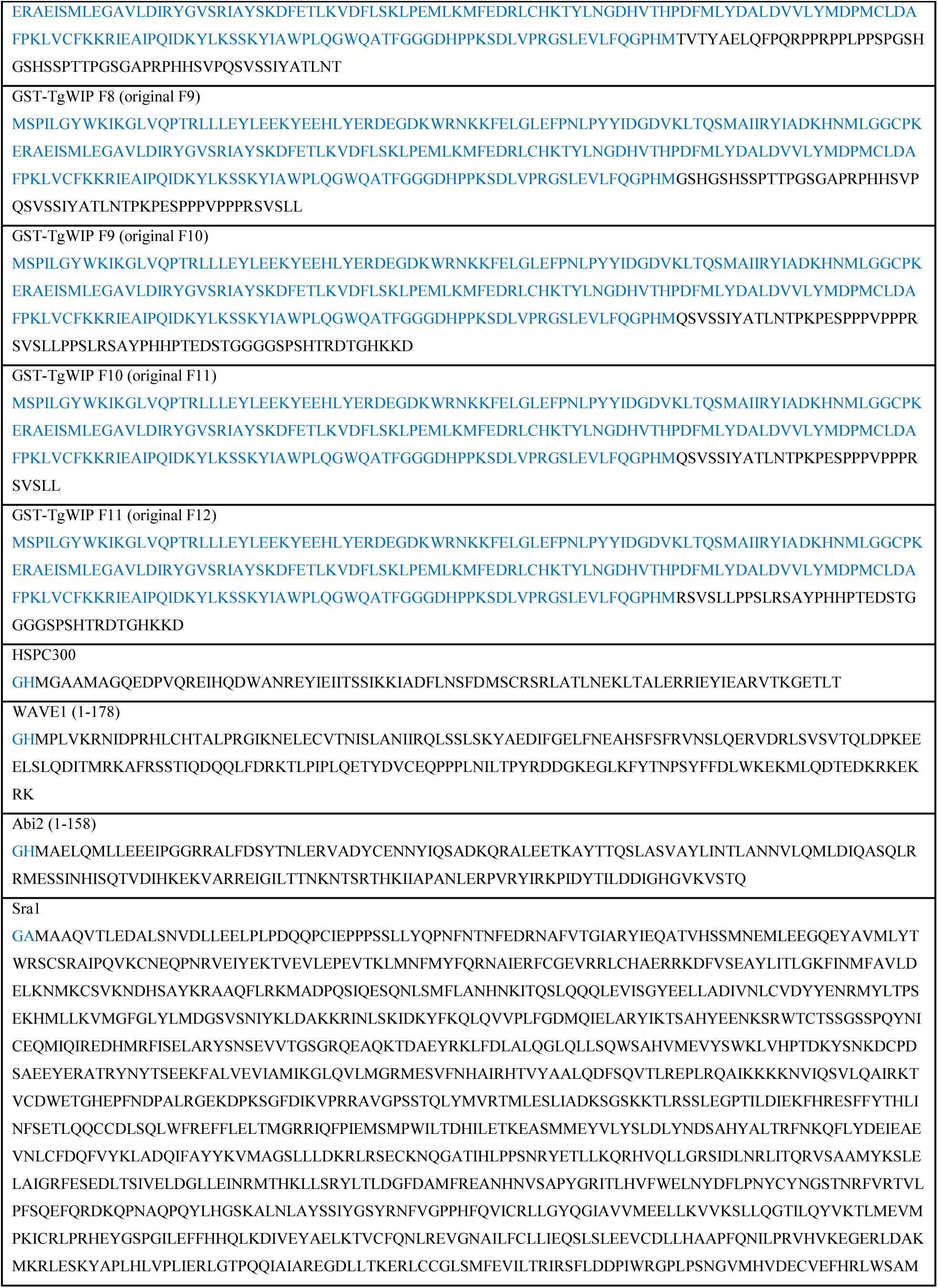
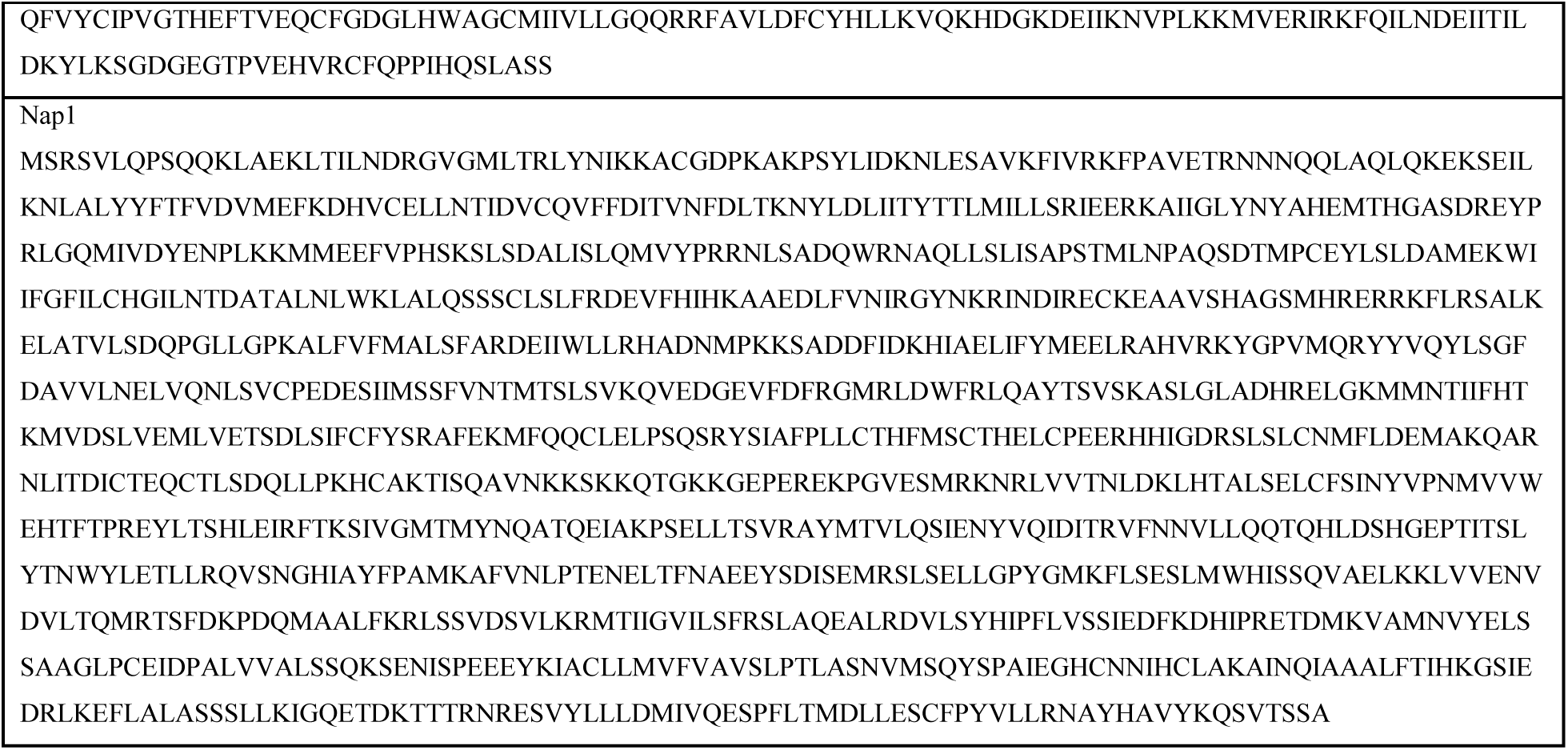
Sequences of recombinant proteins used in this study. Note that only sequences in the final product (i.e., after protease cleavage to remove the affinity tag) are shown and are annotated by corresponding colors.

